# Firefly-inspired vocabulary generator for communication in multi-agent systems

**DOI:** 10.1101/2022.05.19.492736

**Authors:** Chantal Nguyen, Isabella Huang, Orit Peleg

## Abstract

Fireflies’ dazzling light displays are courtship rituals: flying males announce their presence as suitable mates to the females on the ground. Their light signal is composed of a species-specific on/off light sequence repeated periodically. However, thousands of fireflies flashing in a swarm can create immense visual clutter that hinders the detection of potential mates. A partial solution to this visual clutter problem is to flash according to sequences that are more distinct and detectable than those of other individuals. Here, we investigate how distinguishable flash sequences can co-evolve by developing a method for simulating sequences that minimize their mutual similarity with each other while minimizing their energetic cost and predation risk. This simple set of rules produces flash sequences that are remarkably similar to those of real fireflies. In particular, we observe an emergent periodicity in the resulting sequences, despite the lack of any periodicity requirements on the sequences. In addition, we demonstrate a method of reconstructing the evolutionary pressures acting on sets of firefly species. We do so by carrying out simulations that follow known phylogenetic relationships of extant species alongside their characteristic flash patterns.

## Introduction

The multitude of flashes that punctuate a summer’s night are mating calls from fireflies (Lampyridae): a chorus of airborne males announces their presence as suitable mates to females on the ground (Fig. 1A). Fireflies emit light from an abdominal “lantern” organ capable of producing bioluminescence. Some species emit steady glows, while others emit patterns of discrete flashes (Stanger-Hall et al., 2007; Lewis and Cratsley, 2008). Here, we focus on the latter, as these have the potential to temporally encode information and can further provide insight into the development of communications for multi-agent systems (Lewis and Cratsley, 2008).

**Figure 1:**
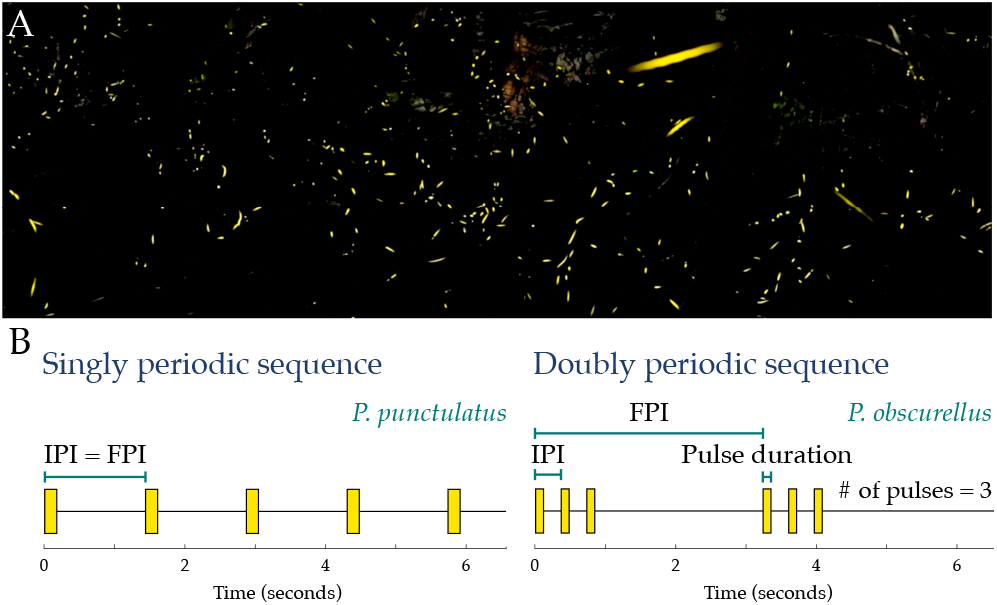
Firefly swarms flash to attract mates. A: Long exposure of a swarm of *Photinus carolinus* fireflies at the Great Smoky Mountains National Park, Tennessee, USA. B: Firefly flash patterns are typically either singly or doubly periodic, as illustrated with the characteristic patterns of *Photinus punctulatus* and *Photinus obscurellus*, respectively. Flash patterns can be parameterized by four values: the pulse duration, the number of pulses in a flash burst, the IPI (interpulse interval, or short period) and the FPI (flash-pattern interval, or long period). For singly periodic sequences, the IPI is necessarily equal to the FPI.

The flying males flash according to a species-specific pattern. Each instance of the pattern is followed by a longer ‘quiet’ or ‘dark’ period wherein the females can respond with a species-specific delay (Lewis and Cratsley, 2008; Stanger-Hall and Lloyd, 2014). If a pair of fireflies recognizes their flashes as indicative of the same species, they can continue to dialogue until they locate each other. On some occasions, fireflies misidentify a signal from a firefly of another species as being from their own. Female fireflies do mistakenly respond to signals from males of the wrong species, while males also locate and approach females of other species, leading to the male realizing the mistake and abandoning the interaction (Lloyd, 1968, 1969; Stanger-Hall and Lloyd, 2014). Moreover, female *Photuris* fireflies, dubbed “femme fatales”, are able to mimic (though not necessarily perfectly) the flash responses of various species of *Photinus* males, luring the male into a deceptive dialogue that results in its attack by the female (Lloyd, 1957; Stanger-Hall and Lloyd, 2014).

While a female’s flash response is relatively simple and often consists of a single flash (Lewis and Cratsley, 2008; Stanger-Hall and Lloyd, 2014), male flash sequences can be more complex. Among North American firefly species, male sequences have been observed to be either singly or doubly periodic (Fig. 1B) (Stanger-Hall and Lloyd, 2014). Singly periodic sequences consist of single flashes (of equal duration) spaced apart by regular quiet intervals. Doubly periodic sequences consist of flash “bursts” of several pulses in a row, separated by longer quiet periods. A flash sequence can be parameterized by 4 values (Fig. 1B): the pulse duration, the interpulse interval (IPI), the flash-pattern interval (FPI), and the number of pulses per pattern. The IPI represents the short period of a sequence, consisting of a pulse and the subsequent pause before the next pulse in a burst. The FPI represents the long period, consisting of the entire flash burst and the succeeding quiet period. If the sequence is singly periodic, the IPI and FPI are equal. As male sequences have been more extensively documented for a range of species than female responses, we focus here on only male signals.

Thousands of fireflies flashing in a swarm presents the possibility of immense visual clutter. This results in a “cocktail party problem”: fireflies must detect the correct pattern of their potential mates while filtering out flashes of other species, similar to how a partygoer can focus on a single conversation while filtering out background music and irrelevant chatter (Bee and Micheyl, 2008). In some species, fireflies synchronize their flashes with swarm mates (Buck, 1988), a social phenomenon that can alleviate visual clutter (Moiseff and Copeland, 2010). The mechanisms, driving factors, and consequences of synchronization in fireflies remain a widely investigated topic (Moiseff and Copeland, 1994, 2020; Sarfati et al., 2021) and source of inspiration for the design of decentralized robotic systems (Christensen et al., 2009; Perez Diaz, 2016).

In more visually complex scenarios, tens of different firefly species can occupy the same geographical area (Lloyd, 1969; Stanger-Hall and Lloyd, 2014). Even if individual fireflies synchronize, it would be beneficial for different species to have distinct flash patterns to facilitate species recognition. Indeed, character displacement has been observed in North American firefly species, where species with overlapping geographical distribution (sympatric) exhibit greater differences in their flash patterns than (allopatric) species that do not (Stanger-Hall and Lloyd, 2014). Sympatric species can differ in their mating times and breeding habitats as well (Lloyd, 1966; Lewis et al., 2004). It has also been observed that fireflies decode species information primarily from the temporal characteristics of flash patterns, namely the pulse duration, number of pulses, IPI, and/or FPI, rather than the color of the light or the movement patterns of the flashing firefly (Lloyd, 1966; Lewis et al., 2004). Hence, we focus solely on temporal information in flash patterns in this paper.

The more conspicuous a flash pattern is, the more noticeable the firefly is to potential mates, but also nearby predators. Bats and spiders, for example, may be attracted to the firefly’s flash (Lloyd, 1973). Predation pressure has resulted in some species foregoing their flash altogether, and instead relying on less efficient pheromone signaling to evade danger (Stanger-Hall et al., 2007; Stanger-Hall and Lloyd, 2014). There may also be energy costs, albeit minimal, associated with flashing (Lewis and Cratsley, 2008), and pressure to minimize energy expenditure would likewise diminish the amount of flashing. As such, we treat both energy cost and predation risk as the same driving force, and will henceforth refer to this cost mechanism as predation risk. Hence, we investigate how firefly flash patterns can co-evolve to be distinguishable under a potentially competing pressure to reduce predation risk.

In this paper, we develop a method for simulating distinguishable firefly-like signals according to an evolutionary process that we term the “vocabulary generator”. In the vocabulary generator, firefly agents possess binary flash sequences; the sequences mutate at a specified rate, and agents adopt others’ sequences in order to minimize a cost function that penalizes both high amounts of flashing (a predation risk), and similarity between sequences of different species. We demonstrate that even when sequences are initialized to random strings of bits that can only change in a bitwise manner, the sequences nevertheless self-organize into near-periodic patterns that resemble real firefly flash sequences. We observe that the competing pressures to minimize predation risk and confusion between species presents a tradeoff, wherein prioritizing discriminability between sequences results in both a higher amount of flashing as well as more complex sequences that contain flash bursts instead of singly periodic flashes. Finally, we explore how the vocabulary generator can be used to “reverse-engineer” a cost function that can capture the relative strength of evolutionary pressures which shaped the flash patterns observed in extant species.

## Vocabulary generator

We introduce the “vocabulary generator”, an evolutionary method for simulating firefly-like signals (Fig. 2). This method is inspired by the naming game procedure, wherein agents achieve common vocabularies via pairwise interactions (Steels, 1997; Baronchelli et al., 2008). In our procedure, multiple firefly species simultaneously minimize a cost function over a series of epochs, whereby fireflies of the same species iteratively compare their sequences via pairwise interactions to mutually adopt the lower-cost sequence. Unlike in the naming game, sequences in vocabulary generator can mutate, as in some evolutionary language models (Nowak and Krakauer, 1999; Nowak et al., 1999). The objective in the vocabulary generator is for fireflies of the same species to arrive at the same characteristic flash sequence, with that sequence being distinguishable from the sequences adopted by other species.

**Figure 2:**
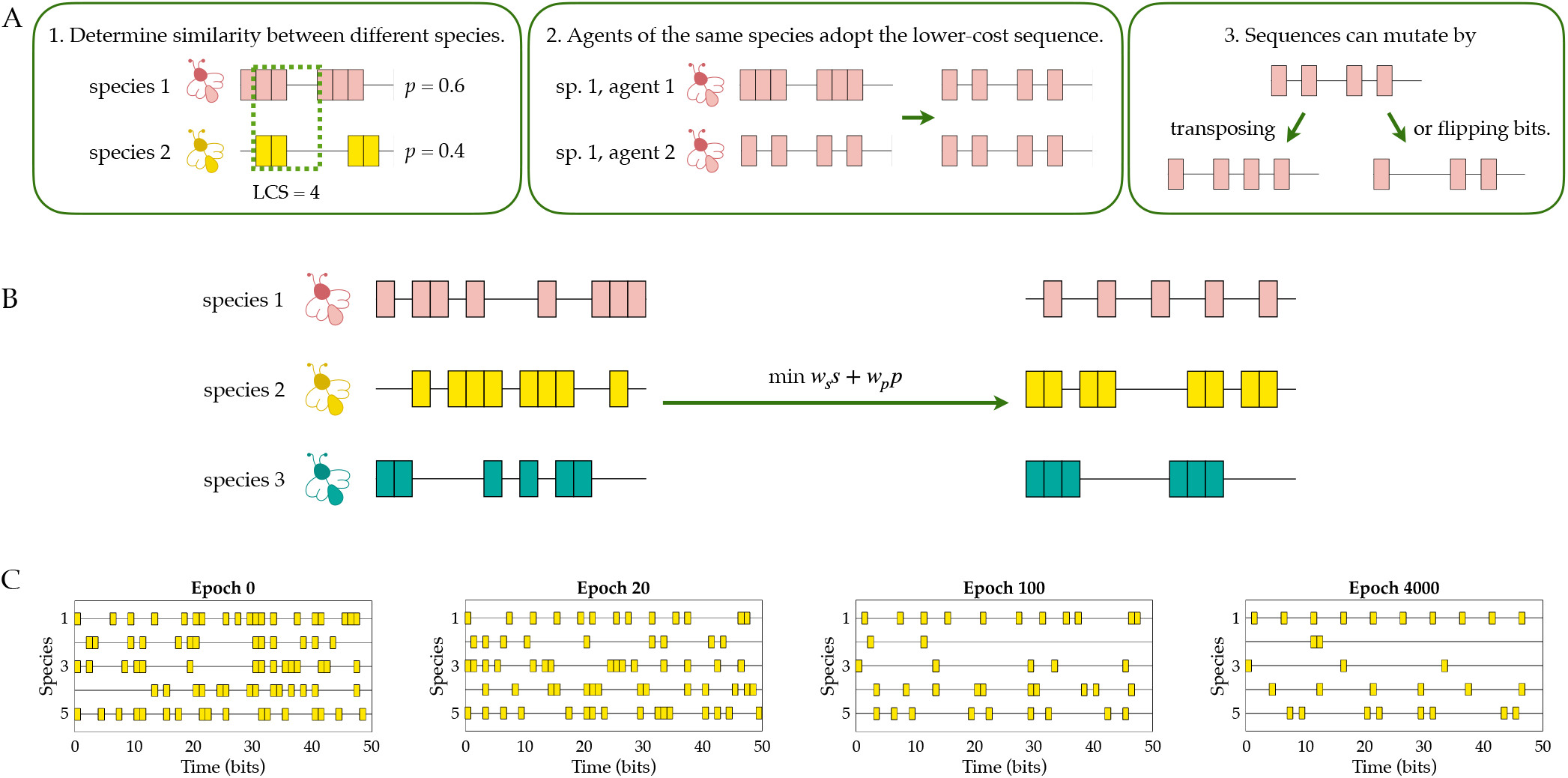
In the vocabulary generator, firefly agents simultaneously minimize a cost function in order to arrive at species-specific flash sequences. A: The similarity between sequences of different species is determined by calculating the longest common substring (*LCS*). In this example, the *LCS* is 4 bits. The predation score *p*, given by the proportion of *on* bits in the sequence, is also shown. Agents of the same species compare sequences, and the lower-cost sequence is adopted by both. Sequences can also mutate by flipping or transposing bits. B: Starting from random initial conditions, multiple species minimize the cost function to achieve distinguishable sequences. C: Evolution of a set of firefly sequences over the course of multiple epochs of the vocabulary generator. Starting from random initial conditions in epoch 0, the five species’ sequences become increasingly periodic by epoch 4000.

A flash sequence of length *L* is defined as a binary string of bits, each with value 1 (*on* or flashing) or 0 (*off* or not flashing):

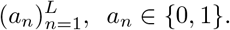

In defining the cost function, we focus on two aspects likely to shape the evolution of firefly communication: the distinguishability between species-specific patterns, and predation risk. We define the cost 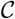 of a particular sequence as follows:

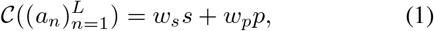

where *s* denotes the average similarity between that sequence and sequences of all other species, and *p* the predation risk of the sequence. *w_s_* and *w_p_* are weights whose relative values can be adjusted to produce sequences with different characteristics.

We define the similarity between two sequences 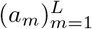 and 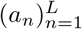 as the length of the longest common substring under cyclic permutation, denoted 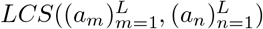, representing the maximal possible overlap between the sequences. We consider all cyclic permutations in comparing the sequences as in nature, there is no guarantee that two different co-habitant species start flashing their sequences at the same time or at a constant offset. The longest common substring between two sequences is given by

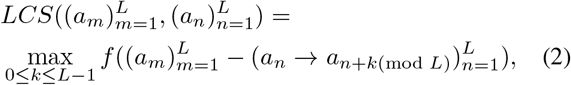

where *a_n_* → *a*_*n*+*k*_ indicates shifting the nth element of the sequence by *k* places, and 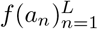 represents a function that computes the length of the longest consecutive subsequence of zeroes in the sequence 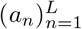. For example, *f* (1, 0, 0, 0, 1, 0, 1) = 3. We note that the spatiotemporal resolution and signal processing mechanisms of real fireflies vary from species to species, and the ability of female fireflies to discriminate and respond to simulated male flashes of varying length or period has been explored for various North American species (Lloyd, 1966; Carlson and Copeland, 1985). These can be further quantified in future field experiments in order to inform the similarity computation. Here we ignore spatial considerations such as the movement of the firefly or the potential attenuation or obscuring of signals by foliage or other obstacles, and we assume that a single bit represents the highest temporal resolution achievable. To compute the similarity term s for a given sequence, the *LCS* between that sequence and all other sequences of different species is averaged and normalized by the length of the sequence, such that s is valued between 0 and 1.

Moreover, we define the predation risk *p* as the proportion of *on* bits over a given sequence length, i.e., 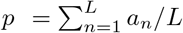. The more a firefly flashes, the higher the predation risk. Likewise, *p* is valued between 0 and 1. Ultimately, this cost function presents a tradeoff: minimizing predation risk will result in sparser flash sequences, but minimizing mutual similarity may require the presence of more flashes to distinguish between species.

In the vocabulary generator (Fig. 2), we define *N_s_* different species of fireflies, with each species containing *N_f_* individual firefly agents. In population genetics simulations, one way to shorten the time to convergence while keeping the mutation rate constant is to introduce the concept of populations for each species (Gillespie, 2004). Hence, we consider a population of *N_f_* agents for each species rather than a single representative phenotype for that species. Each agent is assigned a binary sequence 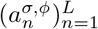, initialized to a random string of bits such that 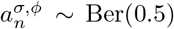, where Ber(*p*) denotes the Bernoulli distribution with probability *p*, *σ* ∈ {1, …, *N_s_*} denotes the species, and *Φ* ∈ {1, …, *N_f_*} denotes the agent identity.

Then, the cost function for a sequence 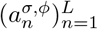 in a system with *N_s_* species and *N_f_* agents per species is given by:

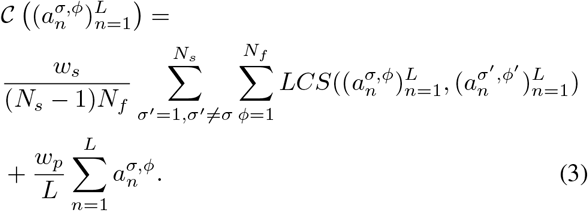

The vocabulary generator method (Procedure 1) is illustrated in Fig. 2A and proceeds as follows. The similarity between sequences of different species is determined by computing the *LCS* (Eq. 2), and the similarity score *s* for each sequence is determined by averaging over the normalized *LCS* between that sequence and those of all other species (Fig. 2A, Box 1). Then, for each pair of firefly agents belonging to the same species, their costs are computed using the previously determined similarity score *s*, following Eq. 3 (Fig. 2A, Box 2). Then, the agent with the higher-cost sequence adopts the sequence of the other agent. The sequence can also mutate with a specified probability (the mutation rate) by flipping a bit (from *off* to *on* or *on* to *off*) or transposing two adjacent bits (Fig. 2A, Box C). The above steps are then repeated until the average costs of the sequences reach a steady state (Fig. 2B). Moreover, Fig. 2C illustrates the evolution of the sequences from a system of 5 species, initialized to random sequences in epoch 0, and reaching a steady state by epoch 4000.

#### Procedure 1 Vocabulary generator

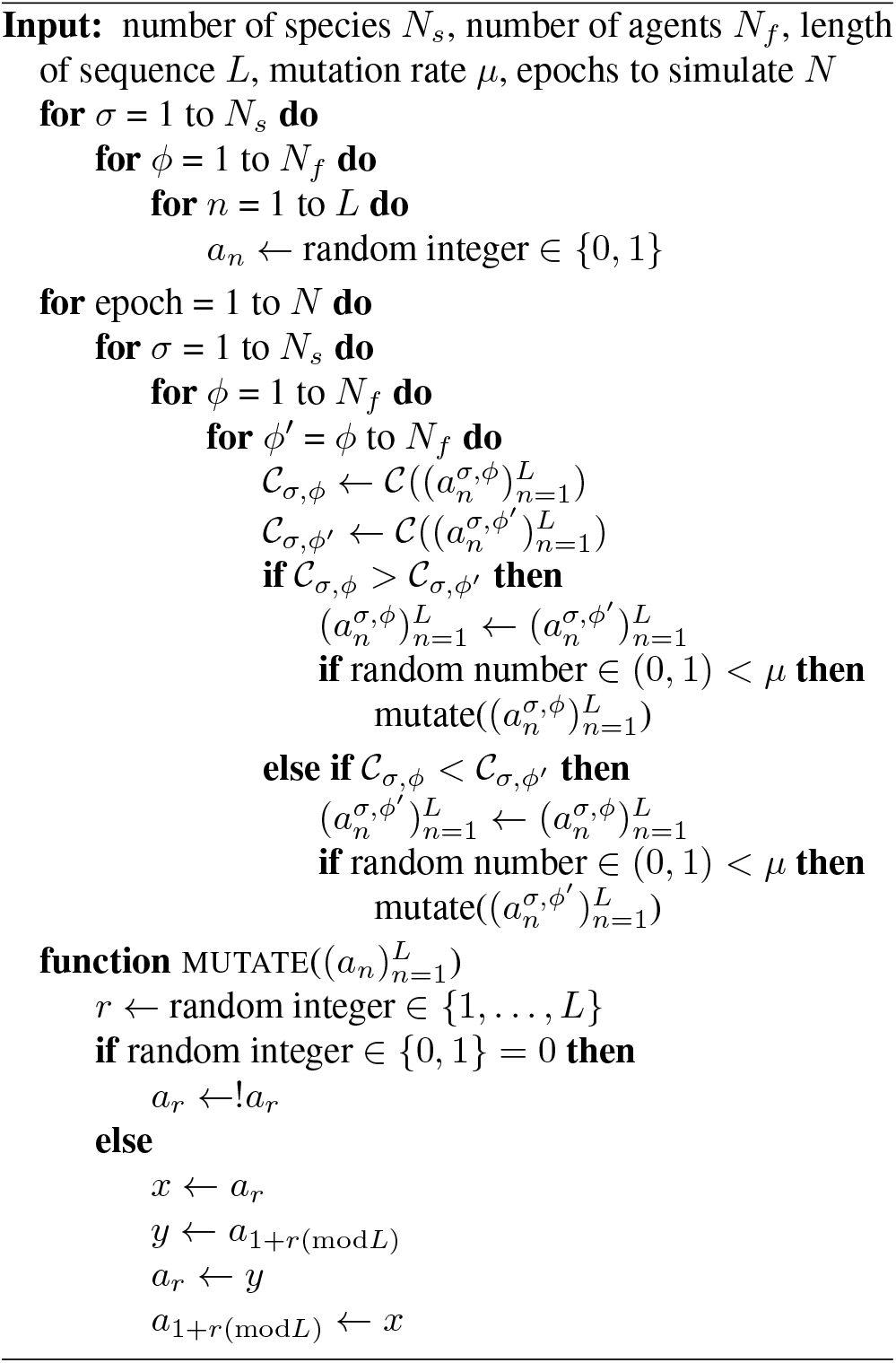

Examples of sequences simulated with the vocabulary generator are illustrated in Fig. 3. The top row shows sequences generated for ensembles of *N_s_* = 7 species with *N_f_* = 10 agents; the resulting characteristic sequence for each species is shown. The middle row illustrates ensembles of *N_s_* = 5 and the bottom row ensembles of *N_s_* = 3, with *N_f_* = 10 agents in both cases. The ratio of the cost function weights *w_s_/w_p_* increases from left to right, with sequences in the left column simulated using *w_s_/w_p_* = 0.4, *w_s_/w_p_* = 1.4 for the middle column, and *w_s_/w_p_* = 2.4 for the right column.

**Figure 3:**
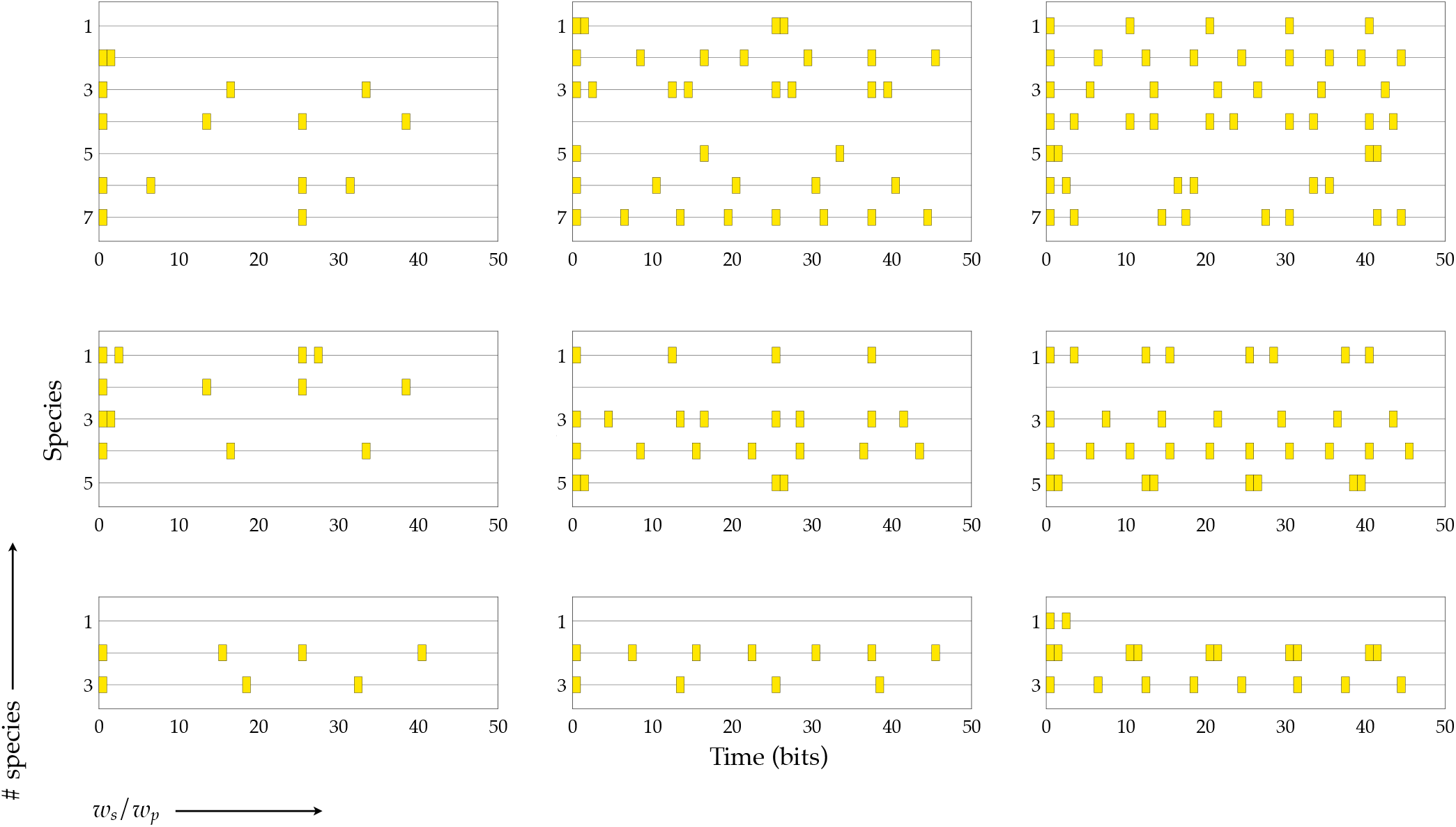
Example ensembles of flash sequences simulated for 3, 5, and 7 species, with weight ratio *w_s_/w_p_* values of 0.4 (left column), 1.4 (middle column), and 2.4 (right column). The weight ratio *w_s_/w_p_* represents the tradeoff between similarity and predation pressures, and higher values of this ratio produce sequences with a greater number of flashes.

We observe that when the *w_s_/w_p_* ratio is small (Fig. 3, left column), sequences are extremely sparse, with more than one sequence being entirely devoid of flashes. As this ratio increases, the frequency of flashing increases, and the length of flashes may increase as well (Fig. 3, right column).

We emphasize that we do not enforce the sequences to be periodic; they are initialized to random strings of bits and mutated in a bit-by-bit manner (Fig. 2A). Nevertheless, we observe that the resulting sequences (Fig. 3) demonstrate an emergent periodicity. We also observe both singly-periodic and doubly-periodic sequences; moreover, we observe that the lengths of individual flashes in a sequence generally do not vary.

We repeatedly run the vocabulary generator to perform a more extensive parameter sweep over values of the *w_s_*/*w_p_* weight ratio that range from 0.2 to 3 in increments of 0.2, and over system sizes ranging between 2 and 7 species (Fig. 4). We quantify the periodicity in the resulting generated sequences by examining the variance in the spaces, or gaps, between flashes. A sequence is classified as singly periodic if the standard deviation of the gap size is less than 0.2 times the mean gap size of that sequence, and doubly periodic if it is greater. Fig. 4A shows the standard deviation in gap sizes as a function of the number of species and the ratio of the similarity and predation weights. For singly periodic sequences, simply the standard deviation of gap sizes, normalized by the mean gap size, is shown. For doubly periodic sequences, the gap sizes are partitioned into two clusters using *k*-means clustering, and the standard deviation for each cluster, normalized to the mean of that cluster, is computed; these two values are then averaged and shown in Fig. 4A. We observe that the higher the *w_s_/w_p_* ratio, the higher the variation in the gap size, and thus the lower the periodicity.

**Figure 4:**
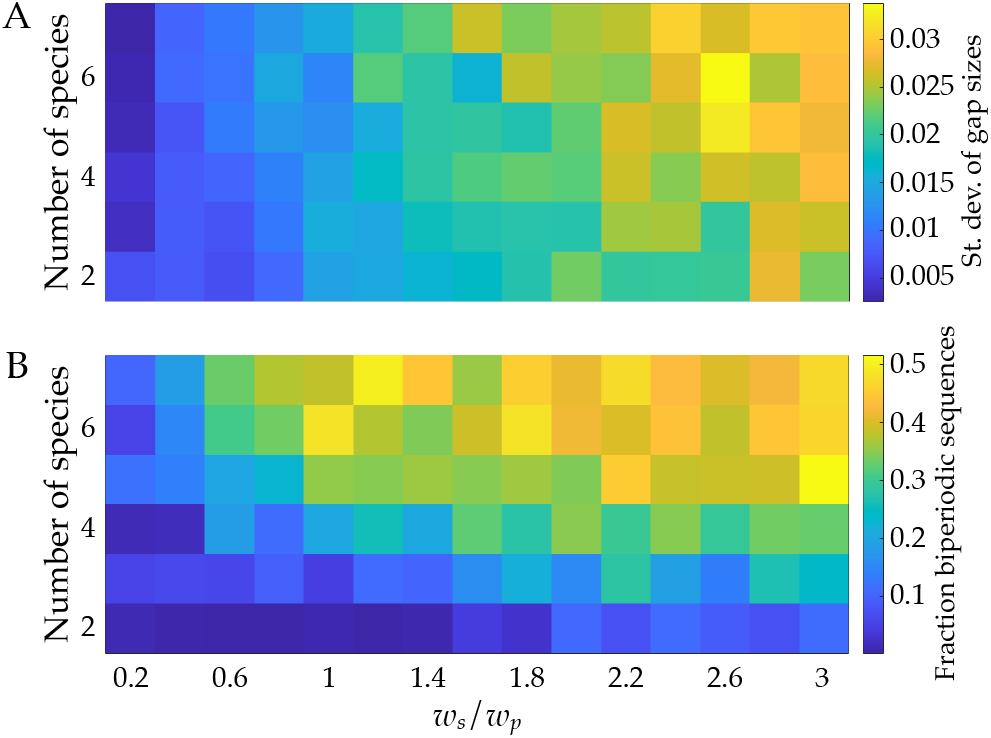
Periodicity in simulated flash sequences. A: Standard deviation in the size of gaps between flashes as a function of the number of species and the ratio of weights *w_s_/w_p_*. The lower the standard deviation in gap size, the higher the periodicity. B: The fraction of doubly periodic sequences present in an ensemble of simulated sequences as a function of the number of species and the ratio of weights *w_s_*/*w_p_*.

The cost function (Eq. 1) captures a tradeoff between predation risk and similarity. When the similarity term is more strongly weighted, i.e., when *w_s_* is larger relative to *w_p_*, we observe that sequences include more flashes in order to differentiate themselves, but also become more complex, containing more sequences with flash bursts. We also observe that for a small number of species, most simulated sequences are singly periodic. For higher numbers of species, along with increased values of the *w_s_/w_p_* ratio, the higher the prevalence of doubly periodic sequences (Fig. 4B). Increasing the number of flashes in a burst can help to reduce the similarity with other sequences when more species are present.

### Brute force validation

To validate our observation of emergent periodicity, we take a brute-force approach by exploring the set of *all* possible binary sequences for a given length. Our objective here is to determine whether sequences that are periodic like firefly flash patterns result in a lower cost (Eq. 1) than those that are not.

First, we generate all sequences of a given length that are unique under cyclic permutation. For example, (1, 0, 1, 0, 0, 0) is equivalent to (0, 1, 0, 1, 0, 0), (0, 0, 1, 0, 1, 0), (0, 0, 0, 1, 0, 1), etc.; and as such only one of these permutations would need to be included. Then, we determine how many of these sequences are valid firefly sequences, i.e., singly or doubly periodic with flashes of equal lengths. For example, (1, 0, 0, 1, 0, 0) is a valid firefly sequence, while (1, 1, 1, 0, 1, 0) is not a valid firefly sequence.

As a specific example, we focus on sequences that are 10 bits long: there are 109 unique binary sequences, and 44 of these are valid firefly sequences (Fig. 5A). Then, from this set of unique sequences, we can generate every combination of 4 sequences representing communities of 4 species, which mimic the ensembles of sequences generated with the vocabulary generator (Fig. 5B). For each of these combinations, we compute the average cost with Eq. 3, with *N_s_* = 4, *N_f_* = 1, *L* = 10, *w_s_* = 1, and *w_p_* = 1.

**Figure 5:**
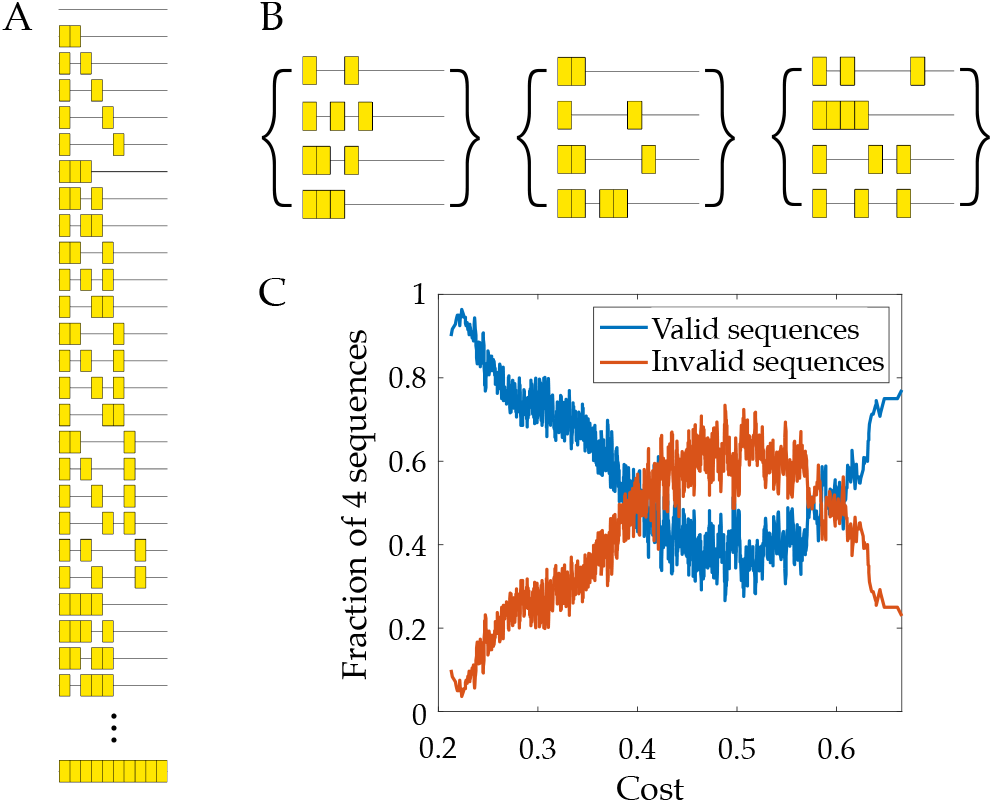
Brute force validation of periodicity. A: 109 10-bit long sequences can be generated which are unique under cyclical permutation; 44 of these sequences are valid firefly sequences, while the rest are invalid as they are neither singly nor doubly periodic. B. Combinations of four sequences are sampled from the set of unique sequences, and the average cost of each combination is computed (Eq. 3, with *N_s_* = 4, *N_f_* = 1, *L* = 10, *w_s_* = 1, and *w_p_* = 1). C. The fractions of valid and invalid sequences in each combination is plotted against the cost of that combination. The blue line (fraction of valid sequences) and orange line (fraction of invalid sequences) sum to a constant value of 1.

We observe that in the combinations with the lowest average cost, a dominant fraction of these are valid sequences (Fig. 5C). However, for combinations with increasing cost, the proportion of invalid sequences increases as well. We note that for very high costs, the majority of sequences in the combinations are valid sequences, but these contain extremely long flashes and are thus energetically costly.

## Reverse-engineering cost functions

Lastly, we demonstrate a way in which the weights *w_s_* and *w_p_* in Eq. 1 can be estimated in order to shed light on how real firefly species may have evolved under selection to minimize similarity and predation risk. Our objective is to use the vocabulary generator to simulate ensembles of sequences for various values of the weight ratio *w_s_/w_p_*, and compare these simulated sequences with known firefly flash patterns to find the weight ratio which results in simulated sequences that most resemble the known patterns. In doing so, this can shed light on the relative importance of the similarity and predation pressures in shaping real firefly signals.

As an example, we consider six species from the Consangineus clade of *Photinus* (Fig. 6A) (Stanger-Hall and Lloyd, 2014). We use the sequences of *P. macdermotti* and *P. ignitus*, which were among the earliest to speciate, as starting “seed sequences” for the vocabulary generator. That is, we initialize a system of *N_s_* = 3 species and *N_f_* agents. The sequences 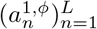 for all *φ* = 1, …, *N_f_* are initialized to all be equivalent to the known sequence for *P. macdermotti*, and likewise 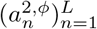 for *P. ignitus*. The sequences 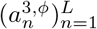 are also all initialized to be that of *P. ignitus*, the most recently speciated species of the two. Then, the vocabulary generator (Procedure 1) is run for a specified number of epochs, where only the sequences 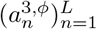 are allowed to change; the seed sequences 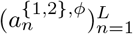 are kept fixed and only factor into the similarity computations (Eq. 2) of the cost function (Eq. 3). By the last epoch, the sequences of species *σ* = *N_s_* = 3 should have all converged to the same sequence, but are assigned to the mode of all 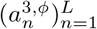 sequences to ensure that they are identical. Then, a simulated speciation occurs: the number of species in the system is increased by 1, so that *N_s_* = 4. The new species’ sequences are initialized to those most recently obtained from the vocabulary generator; that is, 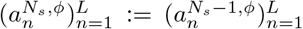, and the sequences 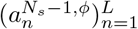 are now kept fixed. The vocabulary generator is then run again, the number of species incremented by 1, and the new sequences initialized to the recent outputs, which are now kept fixed. This is repeated until the desired total number of sequences are obtained, specifically six in our example. The full procedure for simulating sequences based on known phylogeny trees is described in Procedure 2.

**Figure 6:**
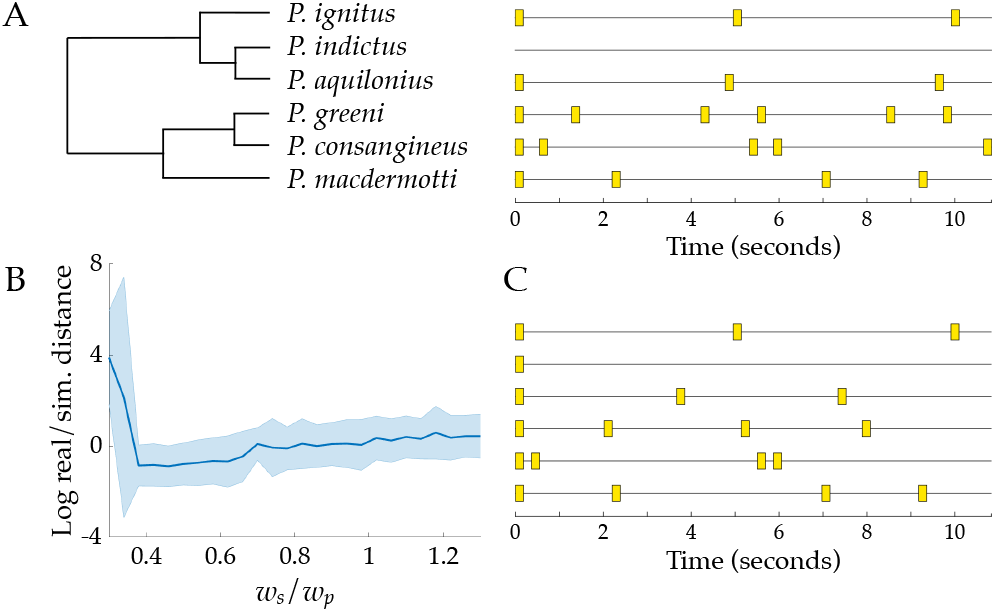
Reverse-engineering the cost function by simulating sequences along the Consangineus clade. A: The phylogenetic relationships of six firefly species and their respective species-specific flash patterns (Stanger-Hall and Lloyd, 2014). B: The logarithm of the average distance between known and simulated flash sequences, as measured by the average root mean squared error in the four flash parameters (number of pulses, pulse duration, IPI, FPI), as a function of the weight ratio *w_s_/w_p_*. Shaded region indicates one standard deviation. C: Examples of simulated flash sequences, obtained following Procedure 2 using *w_s_/w_p_* = 0.46. Note that the first and last sequence are the two “seed” sequences and are identical to the known sequences for *P. ignitus* and *P. macdermotti*, respectively.

#### Procedure 2 Vocabulary generator based on known firefly phylogeny trees

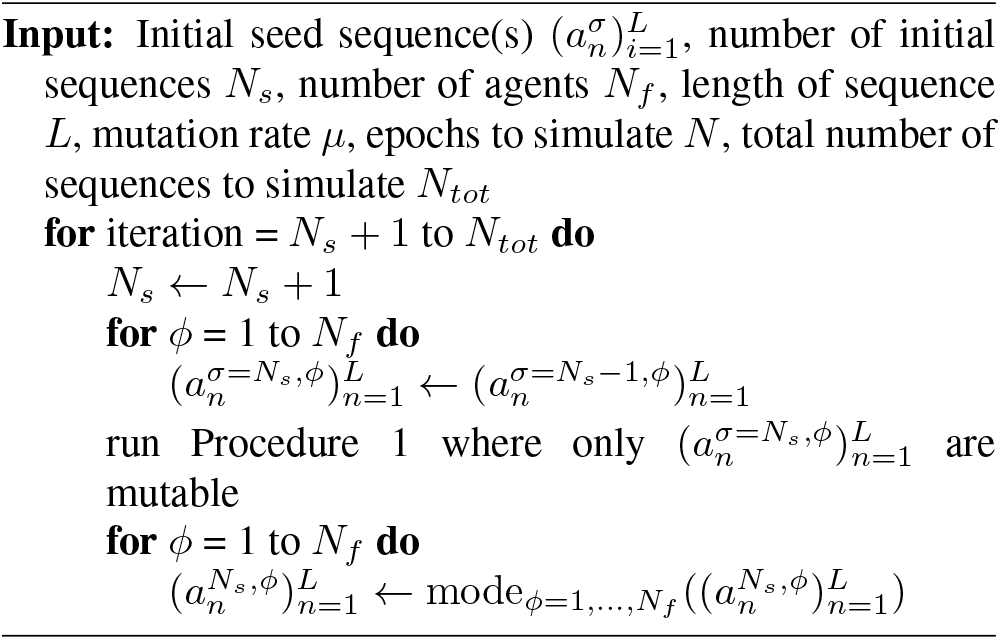

We repeatedly carry out the above procedure (Procedure 2), sweeping over the weight ratio *w_s_/w_p_* in the cost function (Eq. 1) between a range of 0.3 and 1.3 in increments of 0.04. Then, we compare each set of simulated sequences and with the known firefly flash patterns. For each of the known flash patterns, we extract four parameters that describe the sequences: the number of pulses, pulse duration, IPI, and FPI (Fig. 1). We also estimate these parameters for each of the simulated sequences, and compute the average root mean square error between the known and simulated parameters (Fig. 6B). We observe that a weight ratio *w_s_/w_p_* between 0.4 and 0.5 produces flash sequences that are most similar to the known firefly patterns (Fig. 6C).

## Conclusion and Future Work

In this work, we explore how firefly sequences may have coevolved among sympatric species to increase discriminability in order to alleviate the “cocktail party problem” faced by swarms during their flash-mediated mating rituals. We develop a method, termed the vocabulary generator, to simulate the co-evolution of firefly sequences under pressure to minimize both similarity with other species’ signals, and individual predation risk. The resulting simulated sequences are periodic or close to periodic, despite the lack of any constraints pertaining to periodicity. We also observe that combinations of flash sequences that contain periodic sequences can result in a lower average cost than those that contain aperiodic sequences.

We also demonstrate a method in which the vocabulary generator can be used to gauge the relative importance of the selection pressures in the cost function (Eq. 1) in shaping real firefly flash patterns. While we posit here that predation risk and distinguishability play the largest roles in shaping firefly flash patterns, it is possible that other factors may affect communication signals, such as the nature of the habitat and, more significantly, female preference (Lewis and Cratsley, 2008; Stanger-Hall and Lloyd, 2014). Intraspecies variation in male flash patterns has been observed in some species, along with female preference for longer flashes, shorter flashes, or a higher flash rate, depending on the species (Lewis and Cratsley, 2008). Incorporating female responses and the aforementioned factors into the vocabulary generator may shed further insight into firefly signal evolution.

Many other animals, including various species of birds, frogs, and other insects, encode species information in their signals, whether visual or acoustic (Ravignani et al., 2014; Garcia et al., 2020; Höbel and Gerhardt, 2003; Amezquita et al., 2011; Ryan and Rand, 1993). For example, Garcia et al. explored the evolution of species-specific woodpecker drumming patterns and quantified how the mutual information content of the signals changed with species radiation, thereby determining the strength of selection for signal diversity (Garcia et al., 2020). Likewise, in future work, a more extensive analysis of firefly flash patterns could be performed in order to similarly quantify selection pressures on firefly signals, including signal diversity but also predation and other factors, by harnessing the known phylogenetic relationships and recorded signals of numerous North American species and more worldwide (Stanger-Hall et al., 2007; Stanger-Hall and Lloyd, 2014).

In developing the vocabulary generator, we discretize flash sequences into series of bits, where we define a single bit to represent the finest temporal resolution achievable by a firefly’s internal signal processing capabilities. By defining similarity as the longest common substring, we also assume that the species information is encoded as a consecutive sequence of bits. However, fireflies may determine species information by measuring properties relating to the timing of the flash periods or the number of flashes. Moreover, fireflies may also encode species information in non-temporal properties of their signal, such as their movement patterns during flashing or the wavelength of the light, although these properties appear to be less important than temporal characteristics in species identification (Lloyd, 1966; Lewis et al., 2004). Further field experiments can be performed to explicitly quantify the spatial and temporal resolution of a firefly’s signal processing capabilities.

Fireflies, and in particular their synchronization, are increasingly probed as a source of inspiration for swarm robotics (Christensen et al., 2009; Perez Diaz, 2016). Moreover, visible light communication has been explored as a cost-effective method for local, decentralized communication between agents, and firefly signals have also been credited as a source of inspiration in the design of such systems (Maxseiner et al., 2021; Murai et al., 2012; Ito et al., 2018). For instance, an individual robot can be equipped with an LED that can flash according to a pattern similar to that of a firefly’s. We propose that the vocabulary generator method can be adapted to generate different sequences for robotic agents. We note that in the cost function (Eq. 1), predation risk is interchangeable with energetic cost, as the corresponding term serves to minimize the total amount of flashing. For example, for a swarm of robots with differentiated tasks, each group can be programmed to communicate according to its own sequence, optimized to be maximally dissimilar from the others and individually energetically efficient. If the robots must differentiate their tasks in real time, the vocabulary generator could be used to generate easily discriminated sequences rather than using ones that are preprogrammed.

Fireflies have long been a source of wonder and inspiration, but their populations are increasingly threatened by deforestation, urbanization, pesticide use, and climate change (Lewis et al., 2020). Recent research has shown that light pollution can interfere with flash signaling, lowering mating success (Lewis et al., 2020; Firebaugh and Haynes, 2016). Understanding the mechanisms behind firefly communication will be invaluable to worldwide conservation efforts.

## Acknowledgements

O.P. acknowledges internal funds from the BioFrontiers Institute.

